# Comparative evaluation of plasma protein purification and 2-D gel electrophoresis protocols for analysis of HIV-1 infected human plasma proteins

**DOI:** 10.1101/2021.03.03.433830

**Authors:** Sushanta Kumar Barik, Deepika Varshney, Keshar Kunja Mohanty, Deepa Bisht, Shripad A. Patil, Rananjay Singh, Devesh Sharma, SrikanthPrasad Tripathy, Rekha Tandon, Tej Pal Singh, Srikanta Jena

## Abstract

Purification of proteins from human plasma is a herculean task to perform 2-D gel electrophoresis. Human plasma contains nearly 70% albumin and globulin. The removal of such high abundance high molecular weight proteins is very difficult before performing 2-D gel electrophoresis. It becomes more difficult when we intent to investigate in infectious diseases like HIV/AIDS. We tried to the best of our efforts adopting various organic and non-organic based protocols based on various published papers. After failure of these protocols in results of 2-D gel-electrophoresis Aurum serum mini kit (Bio-Rad, USA) was adopted for plasma protein purification for performing 2-D gel electrophoresis. The low-abundance proteins were better resolved by 10% SDS-PAGE in 2-D gel-electrophoresis. Then,we extended the MALDI-TOF/TOF analysis of low-abundance proteins in human plasma by adopting the Aurum serum mini kit (Bio-Rad, USA) for 2-D gel electrophoresis. Thus, we concluded that, depletion of high abundant proteins like albumin and globulin, the use of the Aurum serum mini kit (Bio-Rad, USA) is the protocol of choice to perform the 2-D gel electrophoresis of HIV-1 infected human plasma.

## Introduction

Depletion of highly abundant proteins and enrichment of low-abundance proteins is the first priority in biomarker discovery. Sample preparation by various protocols using magnetic nano particle, chemical depletion and immune-affinity techniques have been used for the depletion of highly abundant proteins in serum. Simple, robust, cost effective, quick and reliable protocols were applied in human serum in the searchfor biomarkers (Jesus et al., 2017). The serum or plasma is the fluid content of the body and contains all the enrichment materials synthesized by the cells. Albumin and globulin are the most abundant proteins in human plasma/serum. The removal of albumin and globulin demands the highest priority during the performance of electrophoresis to search for a biomarker protein in human plasma.TCA/Acetone precipitation method was applied for removal of high -abundance proteins from plasma (Wang et al, 2013).

The presence of large dynamic range of proteins, high abundant proteins, salts, lipids in serum /plasma possesses challenge during the performance of electrophoresis. To overcome such failure conditions in electrophoresis, high-throughput protocols are essential in the field of biomarker discovery(Gong t al, 2008). Several albumin removal methods were compared with specific kits for evaluation of human serum proteome through two-dimensional gel electrophoresis (Fattahi et al, 2012). Other techniques like protein microarrays are essential advance techniques for protein identification in plasma samples. These techniques are useful for the detection of biomarkers in plasma samples (Herosimczyket al, 2006).

The aim of this paper is the comparative analyses of various protein extraction protocols alongwith the protein extraction protocol through the Aurum serum mini kit, Bio-Rad, USA.

## Materials and Methods

### Sample collection

Ten ml of whole blood sample was collected in vacutainer containing EDTA from three healthy individuals at the National JALMA Institute for leprosy and other mycobacterial diseases and three HIV-1 infected patients at ART centre, Sarojini Naidu Medical College, Agra.The plasma was separated by centrifugingat 2000g for 10 min. and stored at-80°C.The viral load was carried out at National Institute for Research in Tuberculosis, ICMR, Chennai, India.

### Chemicals and Reagents

TCA (Trichloro acetic acid, Sigma-Aldrich, USA), Acetone(Mecrk, India), Urea (Sigma-Aldrich, USA), Thio Urea(Sigma-Aldrich, USA), CHAPS(Sigma –Aldrich, USA), ASB-14 (Merck, India), Dithithretol(Sigma-Aldrich, USA), Bromophenol blue (Merck, India), H2O (HPLC purified,Merck, India), Acrylamide(Sigma –Aldrich, USA), Bis-acrylamide(Sigma-Aldrich, USA), Tris base(SRL,India), Hydrochloric acid(Merck, India), Sodium dodecyl sulphate(Merck, India), Ammonium per sulphate(Sigma-Aldrich, USA),TEMED(Sigma-Aldrich, USA),Glycerol(Merck,India), β-Mercaptoethanol(Sigma-Aldrich, USA), Coomassie brilliant blue-R250(Merck, India), Acetic acid(Merck, India), Methanol(Merck,India),-20°C Refrigerator, Centrifuge tube(Sigma G 3-18K), Chloroform(Merck,India), Methanol(Merck, India, Isopropanol (Merck, India), Polyethylene Glycol-6000(Merck, India).

### Protein purification and 2-D gel-electrophoresis

Several methods were developed for the removal of highly abundant proteins from healthy and HIV-1 infected human plasma samples. The purified proteins were quantified through a Bradford assay kit (Sigma -Aldrich, USA). The purified proteins were running in 2-D gel electrophoresis and images were taken and analyzed in gel dock(Bio-Rad, USA). The details of protein purification protocols like TCA/Acetone, PEG precipitation, organic solvents like chloroform and methanol and Aurum serum protein mini kit were followed. Batch wise healthy, and infected purified protein samples were run in standardized iso-electro focusing gradient strips (3-10 pH) and several iso-electro focusing conditions (IEF) were applied to visualize protein spots in the 10% sodium dodecyl sulphate polyacrylamide gel electrophoresis.

### Human plasma protein purification by 50% TCA/Acetone protocol

50% TCA/Acetone protocol was applied to remove highly abundant proteins from HIV-1infected human plasma(Barik et al,2019). The sample was prepared for 2-D gel electrophoresis following passive rehydration for 16-24 h and active rehydration for the following conditions in several steps: step-1. 250V-1hour, step-2. 250V-1. 5hours step-3. 250-3000V-4hours, step-4. 3000V constant until 15Kvh (Bisht et al, 2006).

### Human plasma protein purification by Tri-Chloroacetic Acid /Acetone/polyethylene glycol protocol

The 50% Tri-Chloroacetic Acid /Acetone/polyethylene glycol (16%) was used for protein purification from HIV-1 infected human plasma (Barik et al, 2019). The sample was prepared for 2-D gel electrophoresis following passive rehydration for 16-24 h and active rehydration for the following conditions in several steps: step-1. 0-250V-1hour, step-2. 250V-1. 5hours, step-3 4000-4hours, step-4. 8000-V/hrs (Bisht et al, 2006).

### Human plasma protein purification by organic solvents protocol

Mainly organic solvents were used for protein purification from HIV-1 infected human plasma(Barik et al, 2019).The purified plasma proteins of healthy and infected individuals were run in the modified IEF conditions in several steps:

The sample was prepared for 2-D gel electrophoresis following passive rehydration for 16-24 h and active rehydration for following conditions in seven steps includes, Step-150V -2hr – Linear,Step-2250V -0.15hr –Linear, Step-3 1000V-0.30hr-Slow, Step-4 2500V-0.30hr-Rapid, Step-5 4000V-1.0hr-Rapid, Step-6 4000V for 13,500V/hrs-Rapid, Step-7 500V for 5hrs-slow.

The same organic solvent protocol was used to purify the plasma protein, but different isoelectrofocusing conditions were applied to the same protein purification protocol to get the best spots in 2-D gel electrophoresis. The following modified IEF conditions are given.

### Isoelectrofocusing modified condition-1

A similar protocol was applied to the plasma protein purification and the sample was prepared for 2-D gel electrophoresis following passive rehydration for 16-24 h and certain changes in active rehydration for the following conditions in seven steps. Step-1 50V -2hr – Linear, Step-2 250V-0.15hr –Linear, Step-31000V-0.30hr-Slow,Step-42500V-0.30hr-Rapid,Step-5 4000V-1.0hr-Rapid, Step-6 4000V for 13,500V/hrs-Rapid, Step-7 500V for 5hrs-slow.

### Isoelctrofocusing modified condition-2

The sameorganic solvent protocol was applied for plasma protein purification and the sample was prepared for 2-D gel electrophoresis following passive rehydration for 16-24 h and certain changes in the active rehydration (flow of electric current) for the following conditions in seven steps such as Step-1 50V -2hr – Slow, Step-2 250V -0.15hr –slow, Step-3 1000V-0.30hr-Slow, Step-42500V-0.30hr-Rapid, Step-54000V-1.0hr-Rapid, Step-6 4000V for 13,500V/hrs-Rapid, Step-7 500V for 5hrs-slow.

### Isoelectrofocusing modified condition-3

protein purification from human plasma by organic solvent protocol was applied and the sample was prepared for 2-D gel electrophoresis following passive rehydration for 16-24 h and certain changes in the active rehydration (voltage and flow of current) for following conditions such as Step-1 50V -2hr – Slow, Step-2 250V -0.15hr –slow, Step-3 1000V-0.30hr-Slow, Step-4 2500V-0.30hr-Rapid, Step-5 4000V-1.0hr-Rapid, Step-6 4000V for 10,000V/h-Rapid, Step-7 500V for 5hrs-slow

### Human plasma protein purification by Aurum serum mini kit protocol

The details of the protein purification protocol for human plasma sample are described in the Aurum serum mini kit leaflet and a detailed was followed(BioRad, USA). The sample was prepared for 2-D gel electrophoresis following passive rehydration for 16-24 h. The following active rehydration conditions are as follows :step-1. 50 V - 2hrs-slow, step-2.250 V -0.15hrs-slow,step-3.1000V-0.30hrs-slow,step-4.1500V-0.30hrs-slow,step-5.2500V-0.30hrs-rapid,step-6.4000-3hrs-rapid,step-7.4000-4hrs-rapid,step-8.500-5hrs-rapid.

## Results

The highly abundant plasma proteins were removed from the HIV-1 infected human plasma by the TCA/acetone protocol. Then, the purified sample was electrophoresed in 10% SDS-PAGE. The purified plasma samples wererunning in two-dimensional gel electrophoresis. The 2-D electrophoresis are given in figure-2 and figure-3.

**Fig-1.**
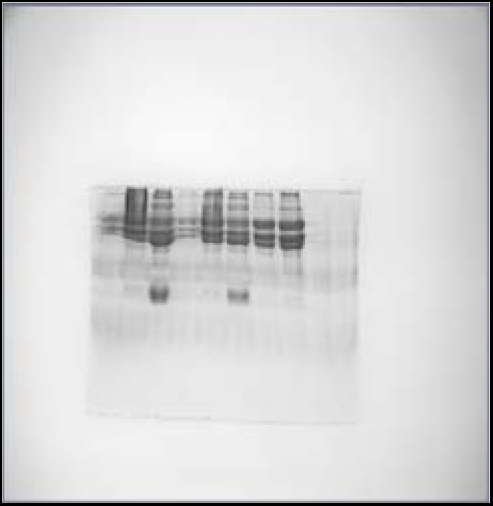
10% SDS-PAGE

**Fig-2.**
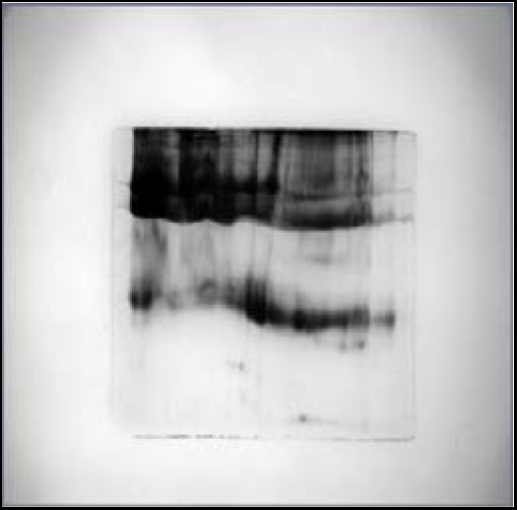
2-D gel electrophoresis

**Fig-3.**
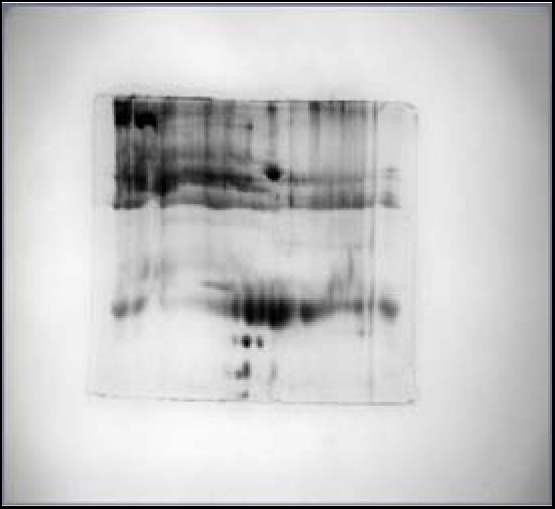
2-D gel electrophoresis

The highly abundant plasma proteins were removed from the HIV-1 infected human plasma by the Tri-Chloroacetic Acid/Acetone/polyethylene glycol protocol. The purified sample was electrophoresed in 10% SDS-PAGE and were running to the two-dimensional gel electrophoresis. The 2-D electrophoresis are given in figure-4-5.

**Fig-5.**
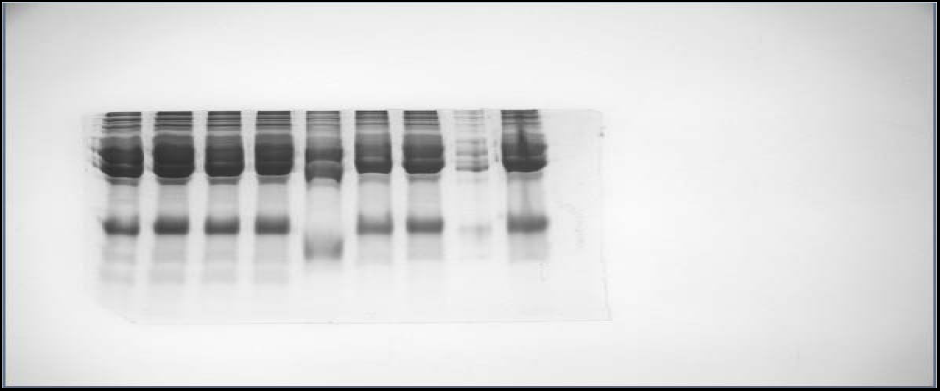
(10% SDS PAGE, Lane-1:4% PEG, Lane-2 :8% PEG,Lane-3 :12% PEG, Lane-4 :16% PEG, Lane- 5: 20% PEG, Lane-6: 24% PEG, Lane-7: 28% PEG).

The organic solvent-based plasma for protein purification protocol was used to remove high abundant proteins from HIV-1 infected human plasma. The 10% SDS PAGE was used to perform one-dimensional and two-dimensional gel electrophoresis. The 10% SDS PAGE and 2-D gel electrophoresis are given in figure -7 and figure-8.

**Fig-6.**
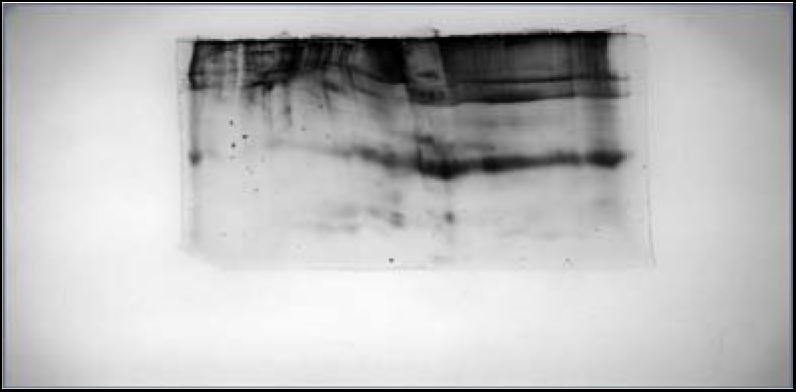
2-D gel electrophoresis

**Fig-7.**
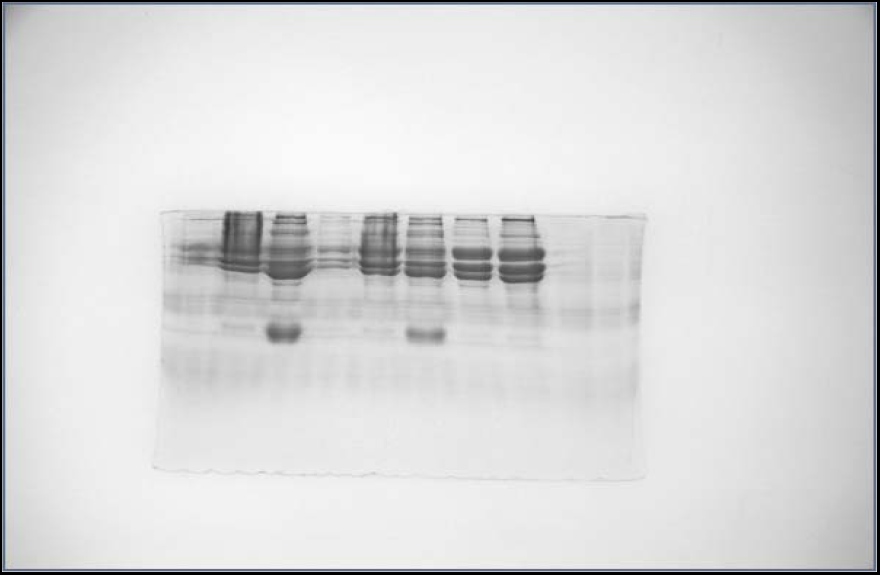
10% SDS-PAGE of extracted protein from human plasma

**Fig-8.**
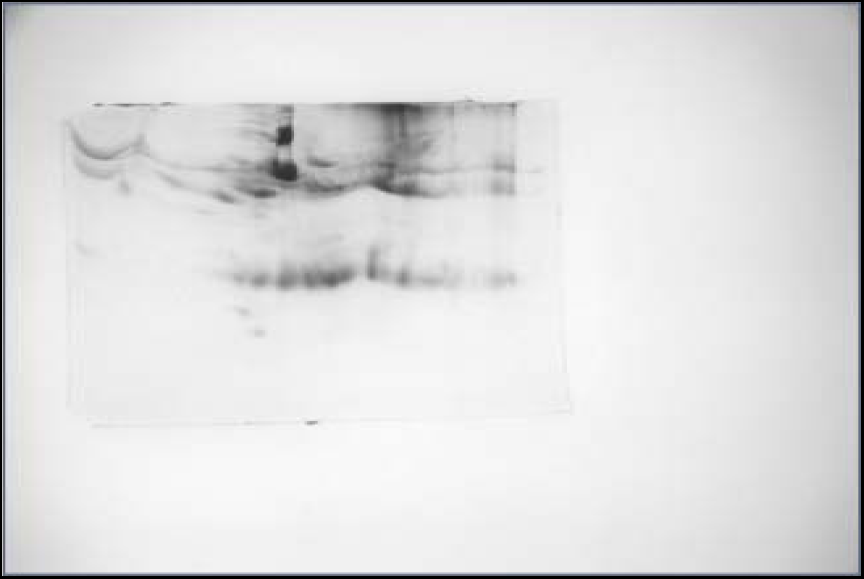
2-D gel electrophoresis

### IEF modified condition-1

The 2-D gel electrophoresis was performed in the IEF modified conditions of the active rehydration step after protein purification by an organic solvent protocol. Figure-9 is given below.

**Fig-9:**
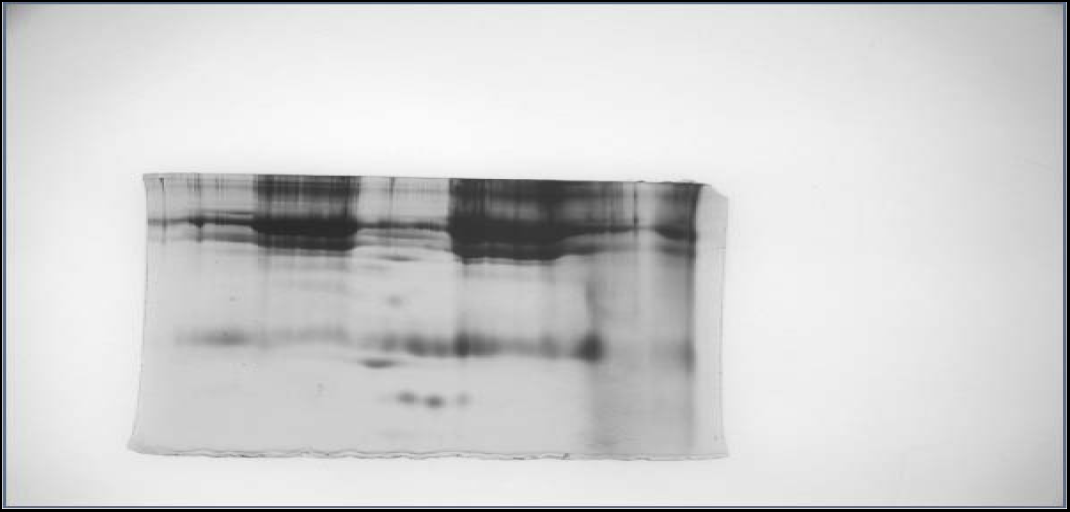
IEF modified condition-1

### IEF modified condition-2

The organic solvent protocol was used to purify the human plasma protein and the modified IEF condition-2 was used to perform 2-D gel electrophoresis. The modified IEF condition figures were given in fig.-10 -11.

**Figure-10:**
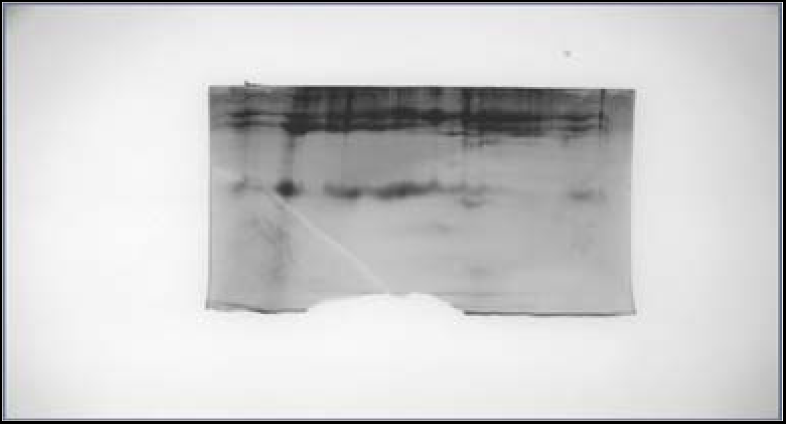
2-D gel electrophoresis

**Fig-11:**
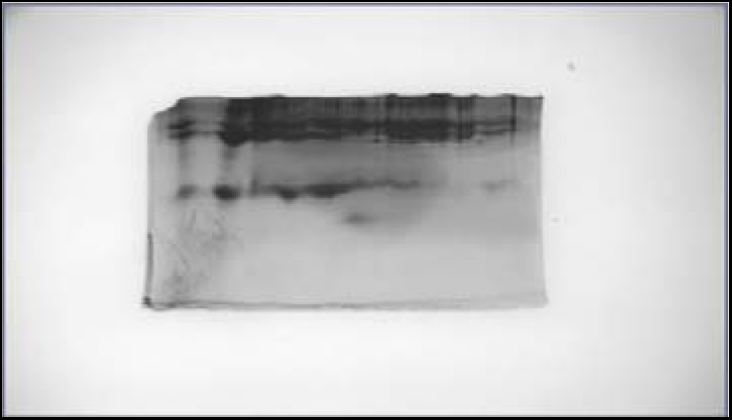
2-D gel electrophoresis

### IEF modified condition-3

The organic solvent protocol was used to purify the human plasma protein. The modified IEF condition-3 was used to perform2-D gel electrophoresis. The modified IEF condition figures were given in fig.-12 -13.

**Fig-12.**
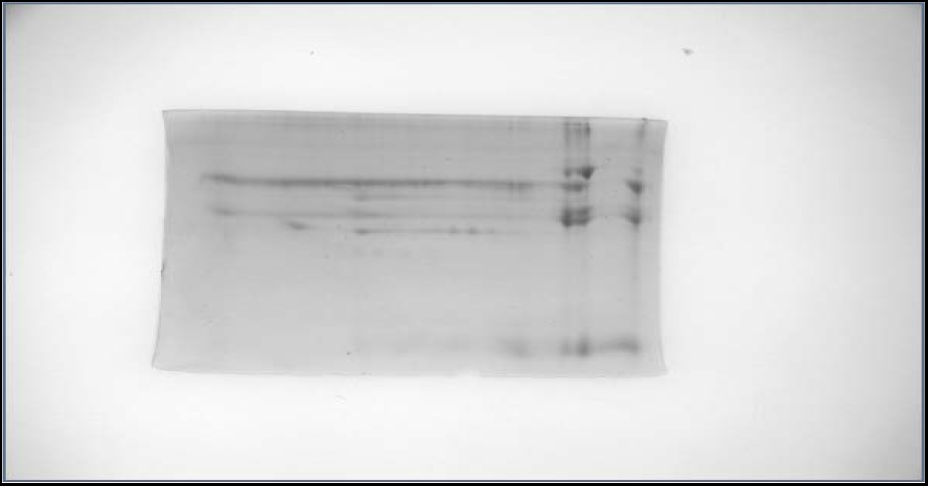
2D gel electrophoresis

**Fig-13:**
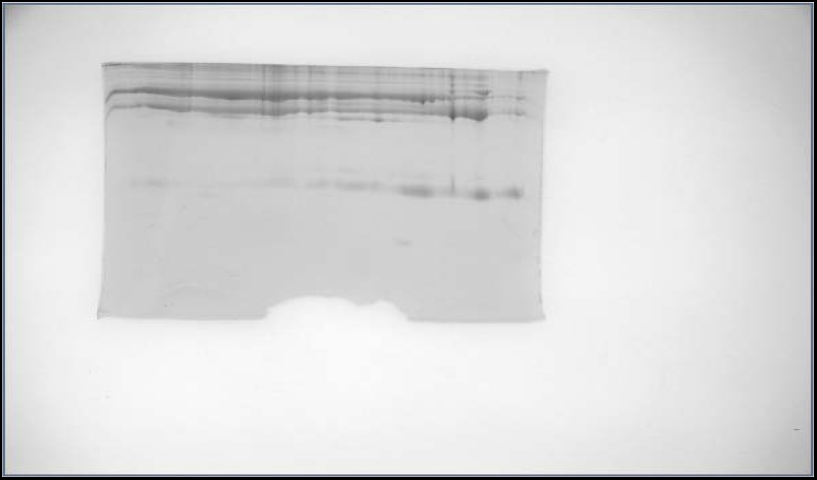
2D gel electrophoresis

The Aurum serum mini kit (Bio-Rad, USA) protocol was used to purify the human plasma protein. The details of the protocol are described in the leaflet. The protocol description is given in supplementary file-1.The plasma protein purified sample was running at 10% SDS - PAGE in one dimensional and two-dimensional gel electrophoresis are given in figure-14 - 15.

**Fig-14:**
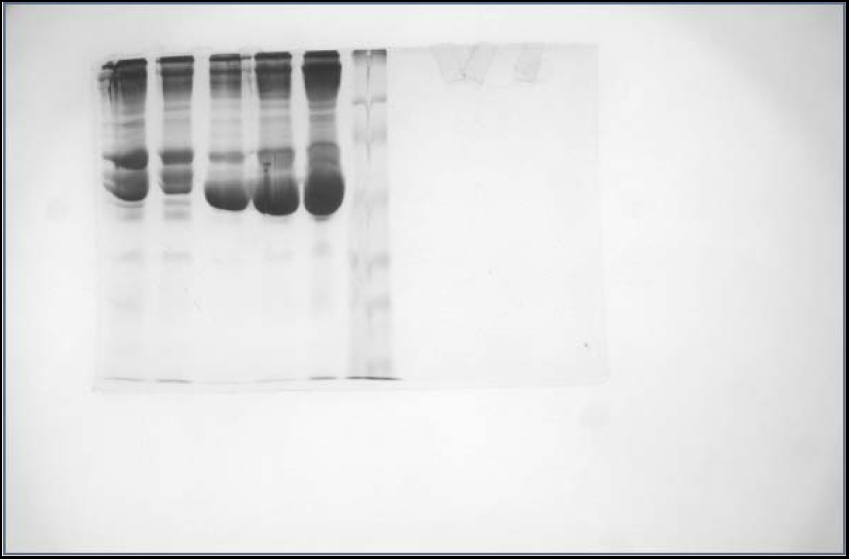
10% SDS -PAGE

**Fig-15:**
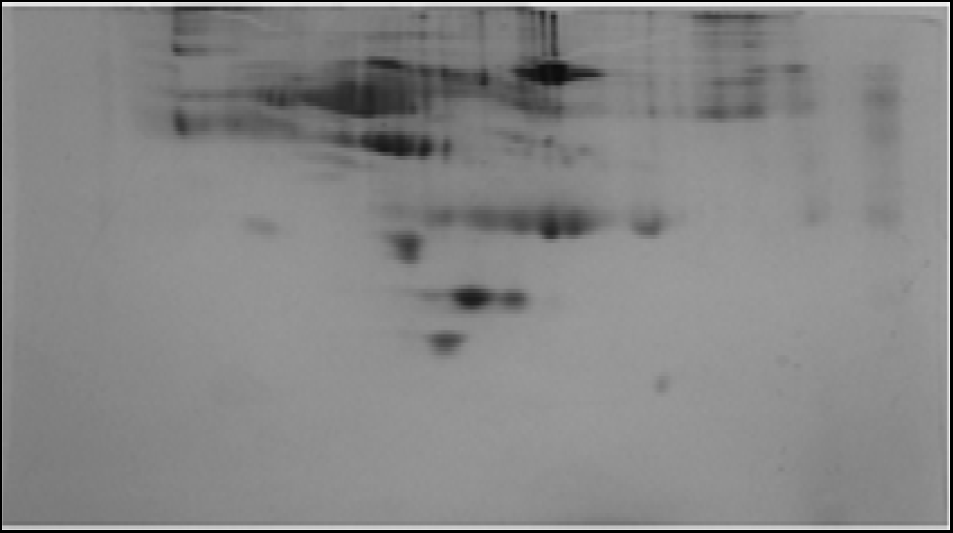
2-D gel electrophoresis

**Table 1.** The comparative results of the various protein purification protocols are given in the table-1.

| SL.No. | Name of protocol | Sample | 2-D electrophoresis results |
| --- | --- | --- | --- |
| 1 | TCA/Acetone | Human plasma | Protein bands are linear, clump and not properly separated in broad range pH 3-10. |
| 2 | TCA/Acetone/PEG | Human plasma | Protein bands are not well separated in broad range pH 3-10. |
| 3 | Organic solvent | Human plasma | Protein spots are not clear to pick up for mass spectroscopic analysis. |
| 4 | Aurum serum mini kit (Bio-Rad, USA) | Human plasma | Protein spots are very clear and in various size, easy to pick up for mass spectroscopic analysis. |

## Discussion

The aim was the comparison of various protocols for the removal of highly abundant proteins from HIV-1 infected human plasma samples and the performance of 2-D electrophoresis. The focus was on a reliable and cost-effective approach to furnishing proteomics experiments. Various protocols were developed to remove the highly abundant proteins and to perform the 2-D gel electrophoresis, but the 2-dimensional gel electrophoresis results were not reproducible except for the kit used for the manufacturer (Bio-Rad, USA)protein purification protocol.

The plasma or serum contains all cellular proteins of patients suitable for thediscovery of the most important biochemical markers. 2-D gel electrophoresis resolved hundreds of thousands of proteinforms in a single analytical run under a good and reliable protocol. Earlier, the 60% TCA/Acetone solution was used for the removal of high abundance, human plasma proteins(Wang et al, 2013). In this plasma proteomics protocol, 50% TCA/Acetone is suitable to remove highly abundant proteins from HIV-1 infected human plasma. Different %concentrations (10% to 80%) of TCA/Acetone were prepared for precipitation of HIV-1 infected human plasma samples and 50% TCA/Acetone was considered for the removal of high -abundance proteins like albumins and globulins in plasma and was resolved by 10% SDS-PAGE. When the purified proteins were 2-dimensional gel electrophoresisunder certain IEF conditions, the protein bands were observed in a linear fashion with improper scattered spots. Simply indicates, the inappropriate use of the TCA/Acetone protocol to perform2-D gel electrophoresis for human plasma samples. Then, we modified the TCA/Acetone protocol with polyethylene glycol-6000 precipitation and the resolution were good in 10% SDS-PAGE. Polyethylene glycol is a water-soluble synthetic polymer played an important role in protein precipitation. The increasing effectiveness of PEG is relatively associated with the polymer size (Polson et al, 1964). In our case, we included the PEG6000 in the TCA/Acetone protocol to improve the detection of low abundance proteins in 2-D polyacrylamide gel electrophoresis. However, the tendency of detection of low-abundance proteins in the human plasma in the 2-D polyacrylamide gel electrophoresis by the iso-electrophoretic condition failed. Thus, our results differ from the Liu et al, 2016 published papers on enhanced detection of low abundance proteins. Michael et al, 1962 adopted a protocol for the isolation of albumin from the blood or human plasma using organic solvents(Michael et al, 1962). To improve, the better resolution of low-abundance proteins in 2-D polyacrylamide gel electrophoresis, the organic solvents were used for human plasma protein purification. Even, we modified several iso-electrophoretic conditions in active rehydration steps, it was promised to fail the detection of low abundance proteins in 2-D gel-electrophoresis organic solvent method. Earlier, a novel and cost-effective method were reported for excess albumin removing plasma for LC-MS/MS analysis of therapeutic proteins(Liu Et al, 2014). Therefore, our organic solvents developed protocol may useful for LC-MS/MS analysis of therapeutic proteins. After the failure of 2-D gel electrophoresis of several protocols developed in our laboratory, then we finally reached the manufacturer protein purification kit Bio-Rad, USA.Then, we first purified two plasma samples by Aurum serum mini kit and then performed 2-D gel electrophoresis in the suitable iso-electrophoretic condition of the active rehydration steps.We found clear, distinguished protein spots in the 10% SDS-PAGE acquainted with the broad range pH gradient strips 3-10.

Electrophoresis separation of proteins always needs a variation of several parameters on published established protocols (Rabilloud et al, 2010).Isoelectric separation of proteins and peptides is an outlook in the field of two-dimensional gel electrophoresis (Pergande et al, 2017). When the classical protocols have reached the limit to validate the new experiments, the search of the new protocol is necessary to get the experimental validation and result oriented objectives by adopting any of one protocol. This article outlines the basics of the isoelectric focusing, summary of the historical achievements of various protocols and the most adoptive one protocol, experimental designs for various isoelectric focusing points and derivative methodologies of the various isoelectric focusing points are also discussed. Various protein purification protocols are also adopted for plasma proteins. This most suitable electrophoretic conditions along with the protein purification protocol could be applicable in the field of biomarker discovery and diagnostic purposes in a disease.

Separation of proteins in blood plasma and serum by two-dimensional gel electrophoresis is most advantaged one (Vesterberg et al,2010). Thus, this study protocol could be utilized in the laboratory diagnosis and clinical investigation of human plasma proteins. High sensitivity analysis of plasma proteins by mass spectrometry is always coupled with iso-electric focusing (Tu et al, 2005). Even we used the broad range of isoelectric focusing gradient strip(IPG, pH 4-7) in this several experimental protocol, the suitable protocol is the outcome in the same IPG and also sensitive for mass spectrometry analysis such as MALDI-TOF-TOF and SWATH-MS. The excellent electrophoretic outcome was obtained during the identification and characterization of serotransferrin and apolipoprotein A1 in HIV-1 patients treated with first line ART (Barik et al, 2020). High resolution two-dimensional gel electrophoresis of plasma protein is essential for identification of known and unknown components in the plasma(Anderson et al, 1977). So, the Bio-Rad protein purification protocol along with the iso-electric focusing point gives the better resolution for identification of specific proteins for mass spectrometry analysis.

Finally, we proposed that this study protocol has already established even the array of electrophoresis protocols are utilized in the field of proteomics.

## Conclusion

Human plasma protein purification for two-dimensional gel-electrophoresis requires standardized and optimized manufacturer protocol as well as the iso-electric focusing point otherwise failure of two-dimensional electrophoresis experiment is remarkable.

## Acknowledgements

ICMR, Govt. of India for providing the ICMR-SRFship to Mr. Sushanta Kumar Barik.

## Supplementary file-1

### Aurum(tm)Serum Protein Mini Kit

#### Protocol Overview

##### Column Setup

1. Place serum protein column in a test tubefor 5 min to allow resin to settle.

2. Remove cap and break tip from column and return to test tube to start gravity flow in column.

3. Wash column with 1 ml of serum protein binding buffer using gravity flow. Repeat.

4. Place column in empty 2.0 ml collection tube and centrifuge for 20 sec at 10,000 x g to dry resin bed. Discard collection tube.

5. Place a yellow column tip on bottom of column and place into a clean 2.0 ml collection tube labeled “unbound”.

##### Sample Binding and Purification

6. In a separate tube, prepare sample by diluting 60 µl serum or plasma with 180 µl of serum protein binding buffer.

7. IAdd 200 µl of diluted serum to top of resin bed in column.

8. IGently vortex column and repeat after 5 and 10 min. Allow column to sit an additional 5 min.

##### Collection of Purified Samples

9. IRemove yellow tip from column and return to tube. Centrifuge column for 20 sec at 10,000 x g, collecting protein fraction in “unbound” collection tube.

10. IUsing same collection tube, wash column with 200 µl of serum protein binding buffer.

11. ICentrifuge column for 20 sec at 10,000 x g, collecting protein fraction in same “unbound” collection tube. Discard serum protein column.

12. The combined fractions contain the albumin and IgG-depleted serum or plasma sample. The sample is now ready for gel analysis.

